# DNA break clustering inside repair domains predicts cell death and mutation frequency in human fibroblasts and in Chinese hamster cells for a 10^3^x range of linear energy transfers

**DOI:** 10.1101/2020.11.30.403717

**Authors:** Eloise Pariset, Ianik Plante, Artem L. Ponomarev, Louise Viger, Trevor Evain, Steve R. Blattnig, Sylvain V. Costes

## Abstract

Cosmic radiation, composed of high charged and energy (HZE) particles, causes cell death and mutations that can subsequently lead to cancers. Radiation-mediated mutations are induced by inter- and intra-chromosomal rearrangements (translocations, deletions, inversions) that are triggered by misrepaired DNA breaks, especially double-strand breaks (DSBs). In this work, we introduce a new model to predict radiation-mediated induction of cell death and mutation in two different cell lines across a large range of linear energy transfer (LET) values, based on the assumption that DSBs cluster into repair domains, as previously suggested by our group. Specifically, we propose that the probabilities of cell survival and cell mutation can be determined from the number of DSBs and the number of pairwise DSB interactions forming radiation-induced foci. We computed the distribution and locations of DSBs with the new simulation code RITCARD (relativistic ion tracks, chromosome aberrations, repair, and damage) and combined them with experimental data from HF19 human fibroblasts and V79 Chinese hamster cells to derive the parameters of our model and expand its predictions to the relative biological effectiveness (RBE) for cell survival and mutation in both cell lines in response to 9 different irradiation particles and energies ranging from 10 to 1,600 MeV/n. Our model generates the correct bell shape of LET dependence for RBE, as well as similar RBE values as experimental data, notably including data that were not used to set the model parameters. Interestingly, our results also suggest that cell orientation (parallel or perpendicular) with respect to the HZE beam can modulate the RBE for both cell death and mutation frequency. Cell orientation effects, if confirmed experimentally, would be another strong piece of evidence for the existence of DNA repair domains and their critical role in interpreting cellular sensitivity to cosmic radiation and hadron therapy.

**AUTHOR SUMMARY:** One of the main hazards of human spaceflight beyond low Earth orbit is space radiation exposure. Galactic cosmic rays (GCRs), in particular their high-charge and high-energy particle component, induce a unique spatial distribution of DNA double strand breaks in the nucleus along their traversal in the cell [1], which result in significantly higher cancer risk than X-rays [2]. To mitigate this hazard, there is a significant need to better understand and predict the effects of cosmic radiation exposure at the cellular level. We have computationally predicted two biological endpoints – cell survival and probability of mutations, critical for cancer induction mechanisms – for the full spectrum of cosmic radiation types and energies, by modeling the distribution of DNA damage locations within the cell nucleus. From experimental results of cell survival and mutation probability in two standard cell lines, we were able to derive the parameters of the model for multiple radiation qualities, both biological endpoints, and two irradiation orientations. The model was validated against biological data and showed high predictive capability on data not used for tuning the model. Overall, this work opens new perspectives to predict multiple responses to cosmic radiation, even with limited experimental data available.

## INTRODUCTION

One of the most concerning health risks associated with long-term missions beyond low Earth orbit is the exposure of astronauts to galactic cosmic rays (GCRs) [3, 4]. With proposed cislunar missions involving the Gateway space station and lunar exploration in addition to upcoming human missions to Mars, GCRs are becoming an increasing concern [5]. However, the health effects of GCRs are poorly understood at this point, because they are primarily extrapolated either from human data following low-linear energy transfer (LET) exposures [6], or from animal and cell work conducted on accelerators simulating acute exposure to some components of a very diverse field of particles [7–9].

The GCRs are primarily composed of protons (85%), helium nuclei (12%), and electrons and positrons (2%) [10, 11]. The remaining 1% of high charge and energy ions (called HZE particles) is the most concerning component of GCRs because of high LET in biological tissues [12], which induces a unique spatial distribution of DNA double strand breaks (DSBs) in the nucleus along the traversal tracks in the cell [1, 13, 14].

One consequence of the elevated ionization power of HZE particles is a relatively higher cancer risk compared to low-LET radiation exposure [2, 15]. Predictions for tumorigenesis induced by HZE are classically scaled linearly from low-LET cancer incidence using the concept of Relative Biological Effectiveness (RBE), defined as the ratio of the dose of a reference radiation (e.g., gamma rays) to the dose of the radiation of interest to produce the same biological effect [16, 17]. The reason for such an approach is that cancer risk in humans has been well characterized for high acute doses of gamma rays (> 0.1 Gy) by monitoring cancer incidence in the cohort of survivors of atomic bombs [18]. Thus, many experimental designs for space radiation studies contain a low-LET component (X-rays or gamma rays) used to scale biological effects towards higher LET, assuming that the same scaling approach can be applied to multiple biological outcomes from low-LET radiation up to high-LET GCRs components. Using this approach, RBE has been used to characterize the impact of HZE particles on different biological responses, such as clonogenic cell survival [19], DNA mutations [20], chromosomal rearrangements [21] and cancer induction [15, 22]. Importantly, RBE evaluations also are being used to develop quality factors in NASA’s cancer risk model [23].

In this work, we focus on two interrelated cellular events important for carcinogenesis: radiation-induced clonogenic death and mutation [2]. These events have been studied previously for V79 Chinese hamster cells and demonstrated distinct LET dependence for mutation and inactivation RBE [24, 25]. Indeed, other *in vitro* studies on mammalian cell survival have reported cell death-based RBE values around 2 following exposure to 1 GeV/n ^56^Fe [26], while cell mutation-based RBE can reach 10 for chromosomal aberrations in similar irradiation conditions, as reviewed by Ritter et al [27].

Our group previously introduced a formalism to predict cell death in a human breast cell line based on experimental data from X-ray irradiation, and expanded it for other LETs using a model of active clustering of DSBs into repair domains, called radiation-induced foci (RIF) [28]. This model offers a computational solution to predict cell death for any other LET by combining the spatial distribution of radiation-induced 53BP1^+^ foci (RIF) with cell survival curves fitted following exposure to X-rays. Here we expand our model to include prediction of mutation frequencies to extend its relevance to cancer risk predictions, by providing a consolidated theory that describes multiple biological endpoints (cell death and mutations) induced by exposure to a wide range of HZE particles.

When we first introduced the active DSBs clustering formalism [28], a simplified microdosimetry profile was used to predict cell death only. In this work, we add more detailed dosimetry profiles using the new simulation code RITCARD (Relativistic Ion Tracks, Chromosome Aberrations, Repair, and Damage) [29] to simulate DSBs in realistic cell geometries. Importantly, we evaluate the effects of the orientation of the irradiation beam in an elongated cell geometry, which is an essential factor in tissues with unique cell geometry (e.g., brain and epithelial cells). While current cancer risk models do not integrate the orientation of irradiation, we demonstrate that it is an important factor to consider to achieve better accuracy of RBE evaluations for cell death and mutation frequency. In summary, we propose a novel unique model that predicts both cell death and mutation frequency for a large range of space radiation-relevant particle types and energies, with LET values ranging from 1 to 2,400 keV/μm. The model is validated against biological data and shows great predictive capability on data not used for tuning the model.

## RESULTS

### Modeling DSBs clustering

As previously introduced by our group [28], we propose to model DSBs clustering using a mathematical formalism that takes into account both the total number of DSBs generated in a cell and the number of pairwise combinations of DSBs inside repair domains. Indeed, the combination of DSB pairs is an essential factor to consider when modeling cell death and mutation induction since it is responsible for increased chromosomal rearrangements. The clustering model based on repair domains has been validated experimentally in human mammary epithelial cells (MCF10A) exposed to X-ray irradiation [28]. This model assumes that the nucleus is divided into separate domains within which all DSBs tend to merge into 1 single RIF [30], with the size of the domains being specific to a cell type and calibrated to experimental data. Figure 1A gives an example of a cell assumed to have 5 domains containing 2 DSBs and 1 domain containing 3 DSBs. The number of pairwise combinations of DSBs (N_comb_) being 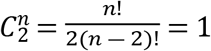 for 2 DSBs and 3 for 3 DSBs, the total number of combinations for the entire cell is: 5 domains × 1 combination + 1 domain × 3 combinations = 8.

**Figure 1.**
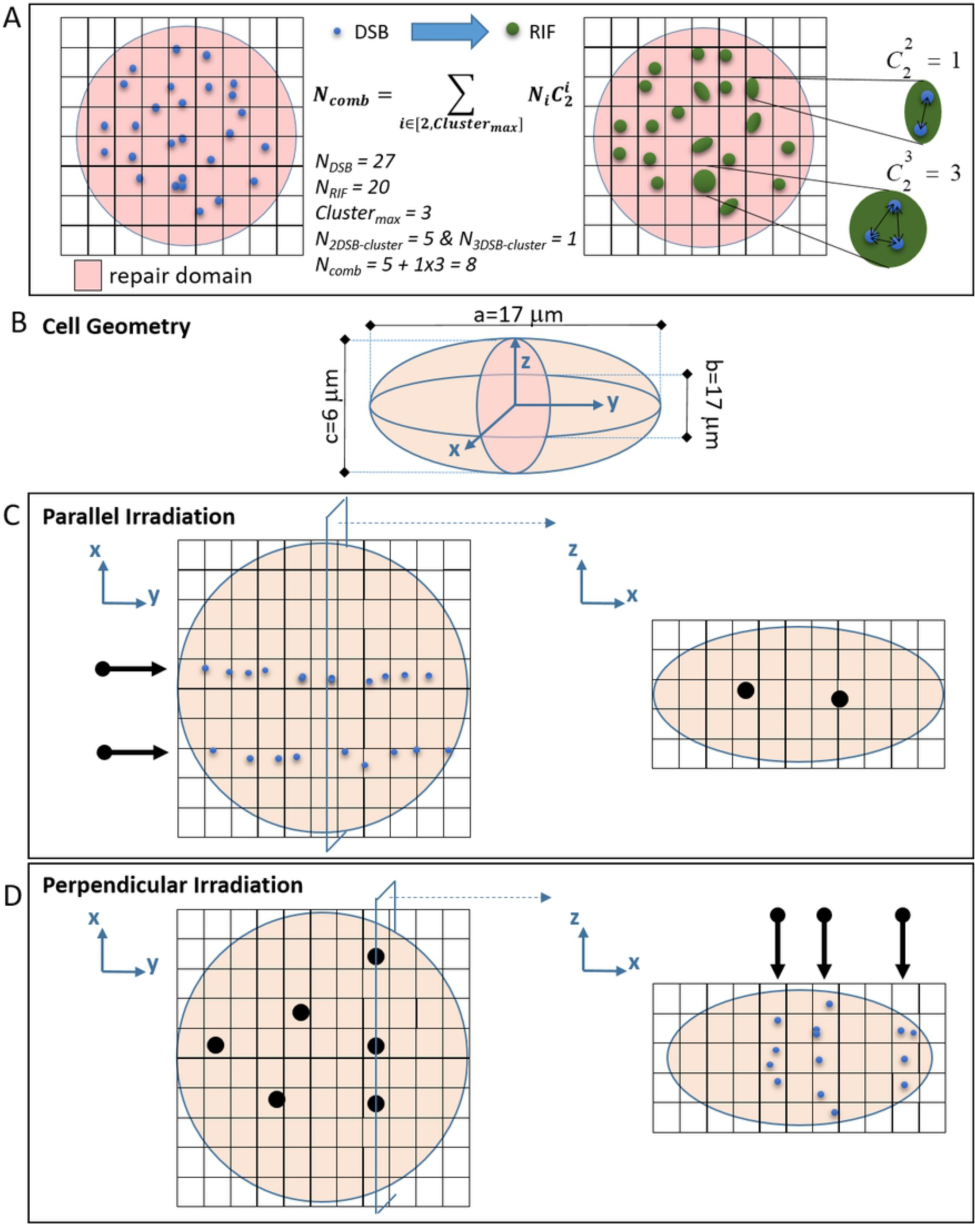
DSBs clustering model, cell geometry and beam orientation. A) Representation of DSB (blue dots) and RIF (green dots) in a cell divided into 60 repair domains (squares). The DSBs within the same repair domain are assumed to cluster into one single RIF. We define the number of DSB combinations in a cell as the sum of all pairwise combinations of DSB in each repair domain. B) 3D representation of the ellipsoid nucleus geometry considered for all simulations. C) Representation of the DSB deposition pattern in parallel orientation, visualized in plane (xz) and (yz). Black dots represent track locations in the irradiated cell, blue dots represent DSB. D) Representation of the DSB deposition pattern in parallel orientation, visualized in plane (xz) and (yz), for the same radiation dose as C).

In this work, we investigate the effect of the cell geometry on DSBs clustering, and subsequently on cell death and mutation frequency. For this reason, we consider the ellipsoid geometry represented on Figure 1B for our simulations of DSBs induction and clustering. The 2 studied irradiation orientations are represented in Figure 1C (beam parallel to the largest axis of the cell) and in Figure 1D (beam perpendicular to the largest axis of the cell). For the parallel orientation, the cross section is shown in the XZ axis view (Figure 1C – dots indicate point of entry of beam), with a smaller area than the perpendicular orientation that has a circular cross section of diameter 17 μm (Figure 1D, XY axis view), typical for both cell type studied: HF19 human fibroblasts and V79 Chinese hamster cells. The number of tracks per cell was computed as described elsewhere [29], with the average number of traversals (λ) given by the fluence of the incident particles (ϕ) and the irradiated area (A): λ=Aϕ. Thus, for a given ion type, particle energy, and dose, the fluence is identical for both irradiation orientations and the number of traversals is proportional to the cross section area of the cell (in our example A(perpendicular)/A(parallel) = (π*17^2^)/(π*6*17) ~2.8, thus 2 traversals in parallel irradiation correspond to 5.6, or about 6 traversals in perpendicular orientation).

### Unraveling dose, LET and orientation effects on DSBs combination and RIF induction

Using the simulation code RITCARD [29], we computed the number of DSBs/Gy, RIF/Gy and DSBs combinations (N_comb_) spanning 10 different sizes of cubical domains, with edge lengths ranging from 0.4 to 2.5 μm, 8 radiation energies (from 10 to 1,600 MeV/n), 4 doses (from 0.5 to 4 Gy), 9 HZE components (^2^H, ^4^He, ^16^C, ^18^O, ^20^Ne, ^28^Si, ^40^Ar, ^48^Ti and ^56^Fe), and 2 beam orientations (perpendicular and parallel) – 1,000 nuclei were simulated for each condition. Figure 2 represents the number of DSBs/Gy, RIF/Gy, and N_comb_ averaged across 1000 simulations for each condition in perpendicular (a) and parallel (b) orientations, for 2 extreme domain sizes simulated (0.4 and 2.25 μm).

**Figure 2.**
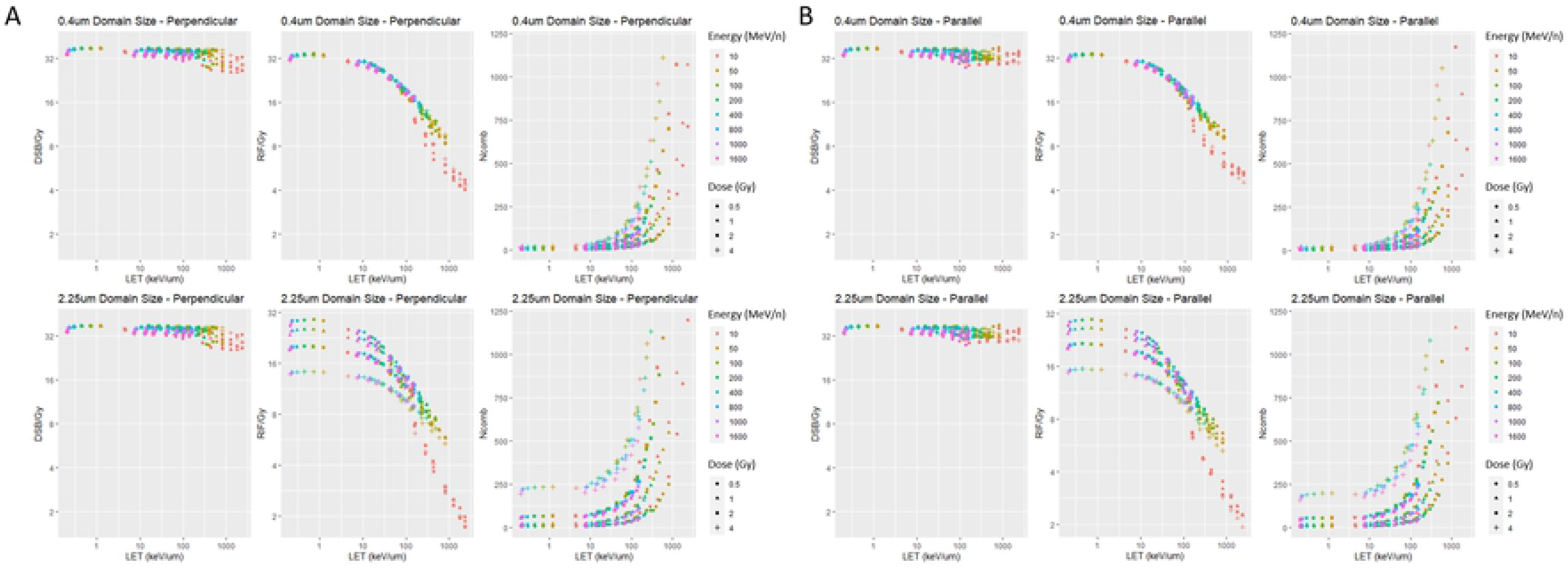
RITCARD global simulation results. Representation of DSB/Gy, RIF/Gy, and N_comb_ averaged across 1,000 simulated nuclei for each irradiation condition, assuming a 0.4 μm or a 2.25 μm domain size, in perpendicular (A) and parallel (B) orientations.

As expected, a DSBs induction yield of about 35 DSBs/Gy is reached for all conditions [31, 32], with a slight drop for LETs above 100 keV/μm, as reported elsewhere [33]. Below 10 keV/μm, the RIF/Gy behavior is dependent on the domain size: 0.4 μm domains show a flat value ~35 RIF/Gy, while for 2.25 μm domains, the plateau drops as the dose increases, reflecting clustering of multiple DSBs into the same domain yielding multiple DSBs/RIF. Above 10 keV/μm, for either domain size, the number of RIF/Gy deviates from the number of DSBs/Gy with a linear decrease following the log_10_ of LET, likely because of an increase in DSBs clustering. Clustering is better visualized by looking at the increasing number of pairwise combinations of DSBs (N_comb_). Thus, below 10 keV/μm, clustering seems to be primarily driven by the dose, with a negligible effect of the radiation energy, while both dose and energy influence clustering at higher LET.

Based on the average values of DSBs/Gy, RIF/Gy, and N_comb_ from 1,000 simulations, no difference is observed between perpendicular (Figure 2A) and parallel (Figure 2B) orientations, suggesting from the averaged results that both orientations lead to similar biological output. However, the distribution of DSBs/Gy (Figure 3A), RIF/Gy (Figure 3B), and N_comb_ (Figure 3C) for all 1,000 simulations in the perpendicular and parallel orientations at both domain sizes (0.4 and 2.25 μm) and for all 4 irradiation doses show slightly different behavior. Such difference results from very distinct average cross section of traversal for the 2 orientations. Briefly, for both domain sizes and beam orientations, we observe a general increase with dose of the mean and the variance of DSBs/Gy, RIF/Gy and N_comb_. While the distribution of DSBs/Gy is independent of the domain size, on average fewer RIF/Gy are obtained for 2.25 μm domains compared to 0.4 μm domains at 1, 2 and 4 Gy, as observed in Figure 2. For a given domain size, the average number of DSBs/Gy is similar for both orientations, but the distribution of DSBs/Gy is slightly larger for parallel compared to perpendicular orientation (Figure 3A), which induces a wider distribution of RIF/Gy as well (Figure 3B). In particular, we notice a higher frequency of cells with low RIF/Gy numbers in the parallel compared to the perpendicular orientation (Figure 3C). As illustrated in Figure 1C and D, a higher number of particles is required for the perpendicular beam to deposit the same dose as the parallel beam, for a given irradiation ion and LET. Thus, in perpendicular orientation, more clustering happens laterally because of shorter inter-track distances, in addition to the longitudinal clustering along the direction of the track, which is similar for both orientations. This might be responsible for a larger number of DSBs per domain, higher clustering and larger RIF that are expected to be more difficult to repair [32], which suggests that the perpendicular orientation could lead to lower cell survival and higher mutation frequencies.

**Figure 3.**
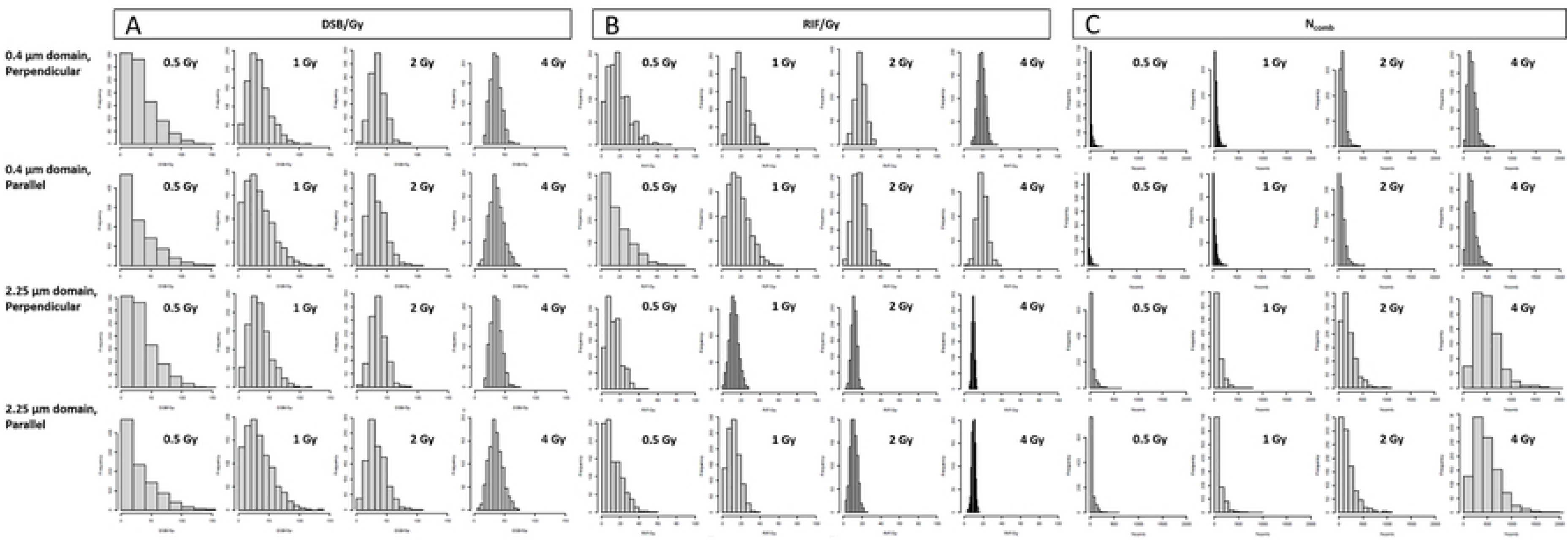
RITCARD individual simulation results. Histograms of the number of DSB/Gy (A), RIF/Gy (B), and N_comb_ (C) at 110 keV/μm (simulation of ^48^Ti irradiation at 800 MeV/n) for 4 irradiation doses (0.5, 1, 2, and 4 Gy), 2 domain sizes (0.4 and 2.25 μm) and 2 beam orientations (perpendicular and parallel). Each histogram corresponds to a simulation of 1,000 nuclei exposed to the same conditions.

### Modeling cell death and mutation frequency from DSBs clustering

As previously introduced by our group [28], we express the survival probability as an average of the individual survival probabilities over all *N_cell_* (= 1,000 in this work) simulated cells, which are obtained from the total number of DSBs generated in each cell (*N_DSB,i_*) and the number of pairwise DSBs interactions (*N_comb,i_*):

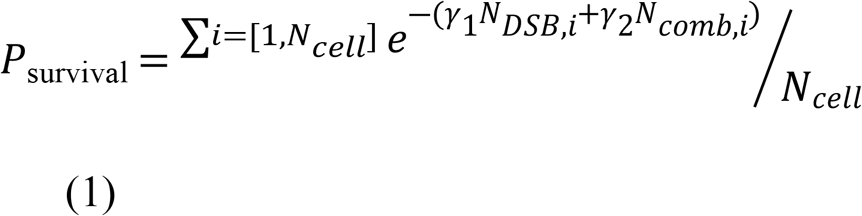

Here we extend our DSBs clustering-based model to the prediction of the probability of mutation, in addition to the survival probability, using the same mathematical formalism. While the classic survival equation involves an exponential dependence on the irradiation dose, the mutation frequency has been commonly described by a linear quadratic equation [24, 34–36]. By extension, we model the probability of mutation in living cells as linearly dependent to *N_DSB,i_* and *N_comb,i_*, weighted by the probability of survival *P_survival,i_* expressed in (1):

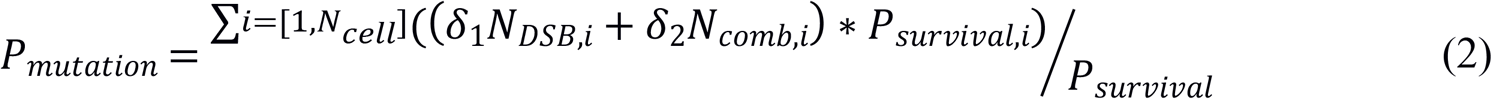

For each of the 1,000 simulated cells, *N_DSB_* and *N_comb_* were predicted by RITCARD simulations for a given repair domain size. The parameters *γ_1_*, *γ_2_*, *δ_1_* and *δ_2_* were optimized to fit experimental data of survival probability in HF19 and V79 cells [34] in the perpendicular orientation. More specifically, all irradiation conditions available experimentally from Cox et al. [34] were tested in RITCARD for 10 different domain sizes between 0.4 and 2.5 μm: X-rays, ^4^He at 20, 28, 50, 70, and 90 keV/μm and ^11^B at 110, 160, and 200 keV/μm for HF19; X-rays, ^4^He at 20, 50, and 90 keV/μm and ^11^B at 110 and 200 keV/μm for V79. Note that as a first approximation the same cell geometry was used for simulations of HF19 and V79 (Figure 1B). For each domain size and LET condition, 4 irradiation doses were tested (0.5, 1, 2, and 4 Gy). The optimal parameters *γ_1_* and *γ_2_* were chosen to minimize the residual across all 4 doses between the experimental survival probability reported in [34] and the computed survival probability obtained from simulations of *N_DSBs_* and *N_comb_*, for each cell type, domain size, and LET condition. Similarly, the parameters *δ_1_* and *δ_2_* were optimized to fit the computed frequency of mutation to experimental data reported in [34]. The computed curves for survival and mutation using the optimal parameters for each condition are represented in Figure S1, and compared to experimental values (black points). Based on this calibration of our model, we identified the domain size that provides the best fit to experimental data across all LET conditions, and for both cell types: a domain size of 2.25 μm was considered for the rest of this work.

Interestingly, the parameters optimizing our model for a 2.25 μm domain were dependent on the LET (Figure 4).

**Figure 4.**
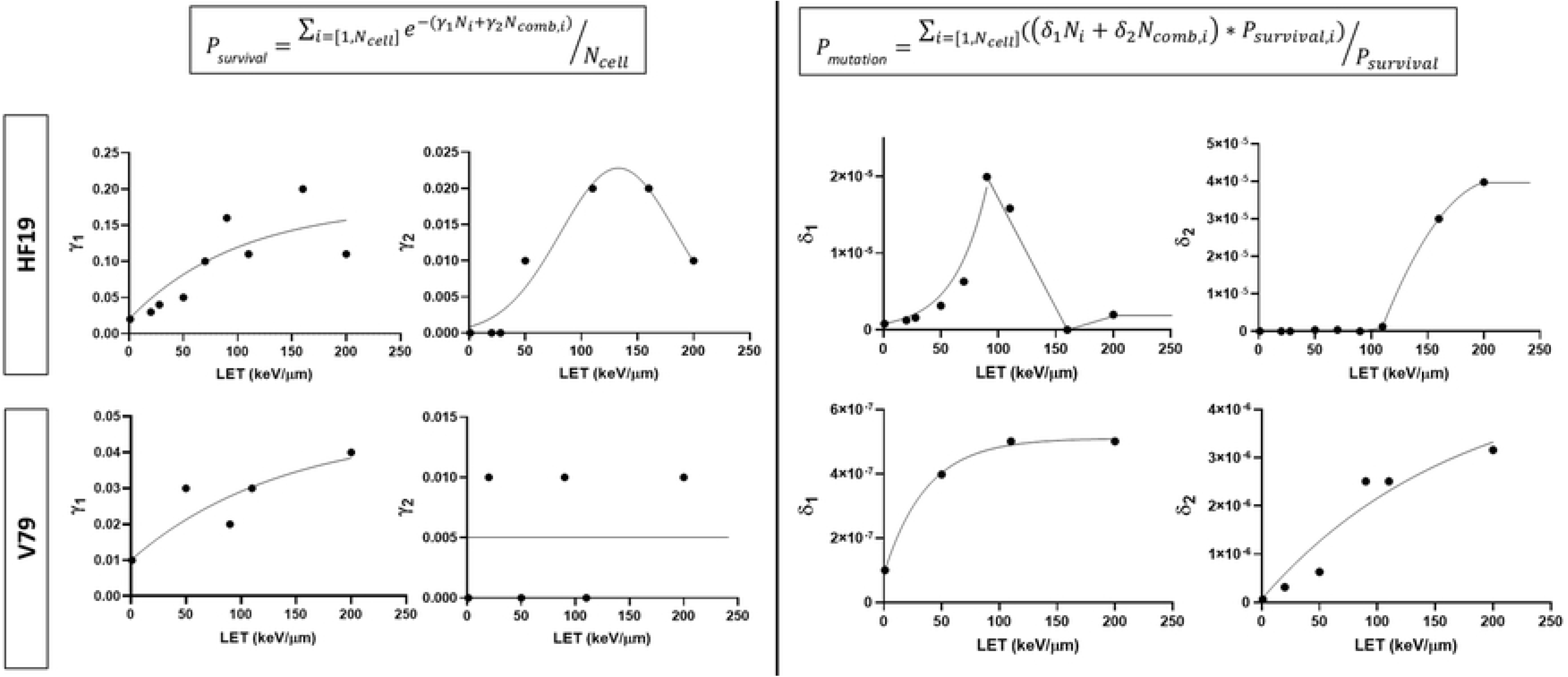
Parameters of the models for survival probability and mutation frequency. Representation of the optimized parameters (*γ*_1_ and *γ*_2_ for survival, *δ*_1_ and *δ*_2_ for mutation) obtained from best fit to experimental data of survival probability and mutation frequency in HF19 and V79 at each reported LET (dots) [34], and derived models (solid lines).

Thus, the following functions were used to model the LET-dependence of all 4 parameters:

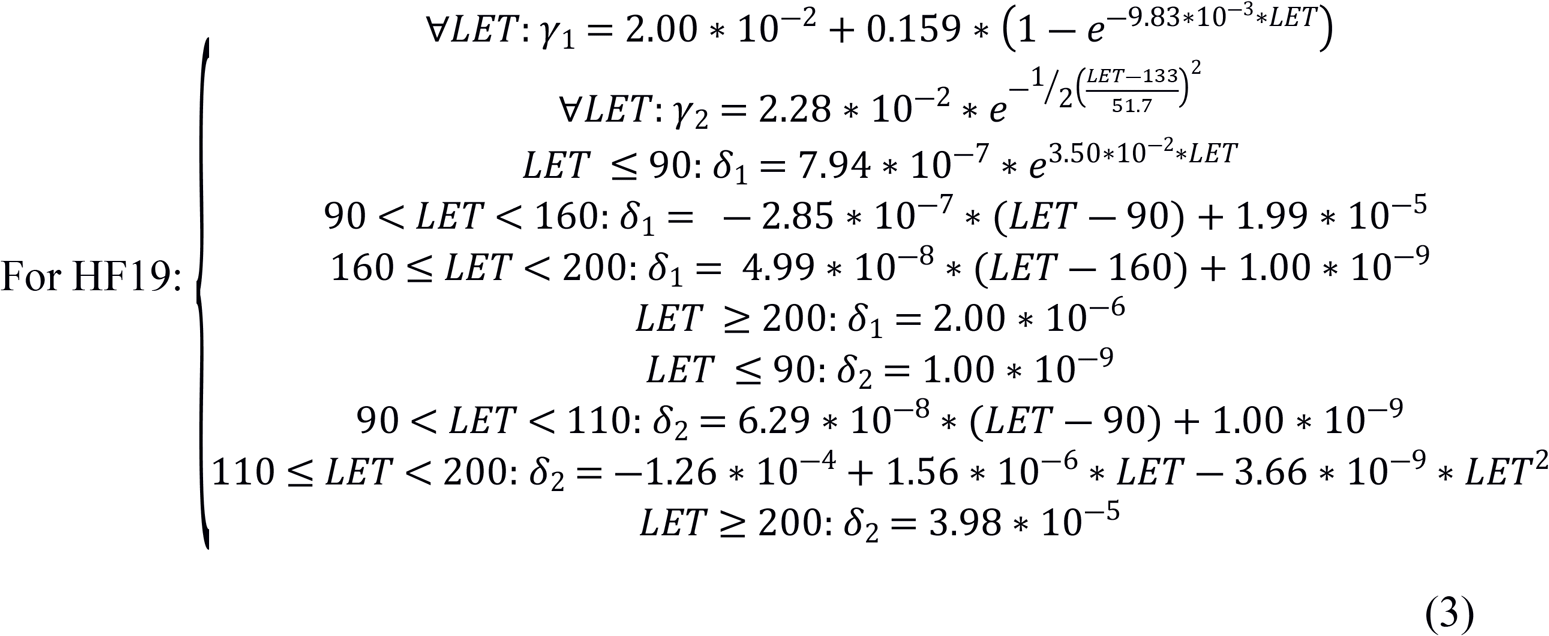

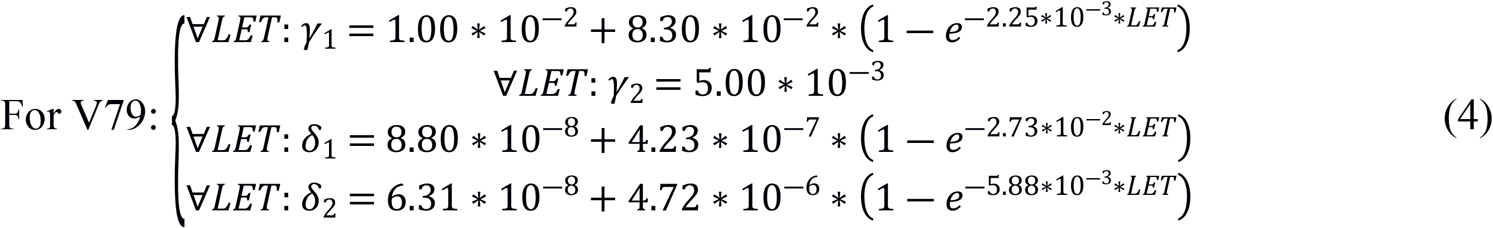

These parameters were used to extend our predictions for survival and mutation frequency probabilities to any other irradiation conditions applied on HF19 and V79 cells. However, because of the LET dependence of the parameters, our model has to be extrapolated when considering LETs outside of the range of irradiation conditions available for calibration.

### Predicting RBE for cell death and mutation frequency

Using the code RITCARD with a domain size of 2.25 μm, we generated 1,000 nuclei for a total of 576 simulation conditions (^2^H, ^4^He, ^16^C, ^18^O, ^20^Ne, ^28^Si, ^40^Ar, ^48^Ti, ^56^Fe; kinetic energy E = 10, 50, 100, 200, 400, 800, 1000, 1600 MeV/n; dose D = 0.5, 1, 2, 4 Gy; perpendicular and parallel orientations). This led to a large coverage of LET from ~0.2 keV/μm to ~2300 keV/μm. Note that above 200 keV/μm, no experimental data were available to calibrate the model parameters, and extrapolations were used, as described in equations (3) and (4) (the data point for Nitrogen irradiation at 470 keV/μm was not used for calibration due to negative fitting parameters reported by Cox et al. [34]). For each simulation condition, probabilities of survival and mutation frequency were derived from the computed *N* and *N_comb_* using equations (1) and (2), respectively. Based on these simulations of predicted survival and mutation frequency, a fit was applied to extract the classic coefficients α and β of the survival and mutation equations, using a least square fit method:

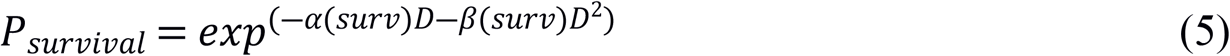

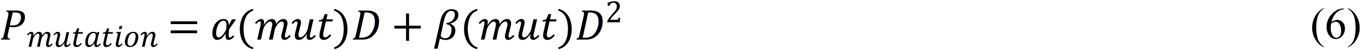

The results of our simulations and respective fits are represented for survival (Figure 5A for HF19, Figure 6A for V79) and mutation (Figure 7A for HF19, Figure 8A for V79) for all irradiation conditions, in parallel (red) and perpendicular (blue) orientations, and compared to experimental X-ray results (black), reported by Cox et al. [34]. From these results, we report RBE for survival and mutation frequency based on α and β values, using ratios to the experimental α and β values for X-ray irradiation [34]. Note that for HF19 cells, β was set to zero for both cell death and mutation in the paper we used as an experimental reference [34], so we applied the same condition in our model. For both cell types, we observe the classic peak of RBE around 200 keV/μm for survival (Figures 5B and 6B) and mutation (Figures 7B and 8B) RBE_α_, followed by an overkill effect at higher LETs [28, 37–39] for which an excess of energy is deposited by the particle to induce cell death or mutations, thus inducing a lower effect per absorbed dose. The respective peaks of RBE_α_ are: 4.6 (perpendicular) and 3.4 (parallel) for HF19 survival (to compare to 4.0 in calibration data), 18.4 (perpendicular) and 6.3 (parallel) for HF19 mutation (to compare to 7.1 in calibration data), 9.5 (perpendicular) and 7.2 (parallel) for V79 survival (to compare to 9.0 in calibration data), 37.8 (perpendicular) and 28.0 (parallel) for V79 mutation (to compare to 18 in calibration data), the error on these values being determined by the models defined in Figure 4 for the dependency of the fitting parameters with LET. Notably, our model also predicts experimental data that were not used for calibration, with RBE values for mutation reported to reach up to 23.7 at 110 keV/μm in V79 cells [40], which matches even better the predicted peak for RBE_α_ than the data used for calibration (Figure 8B).

**Figure 5.**
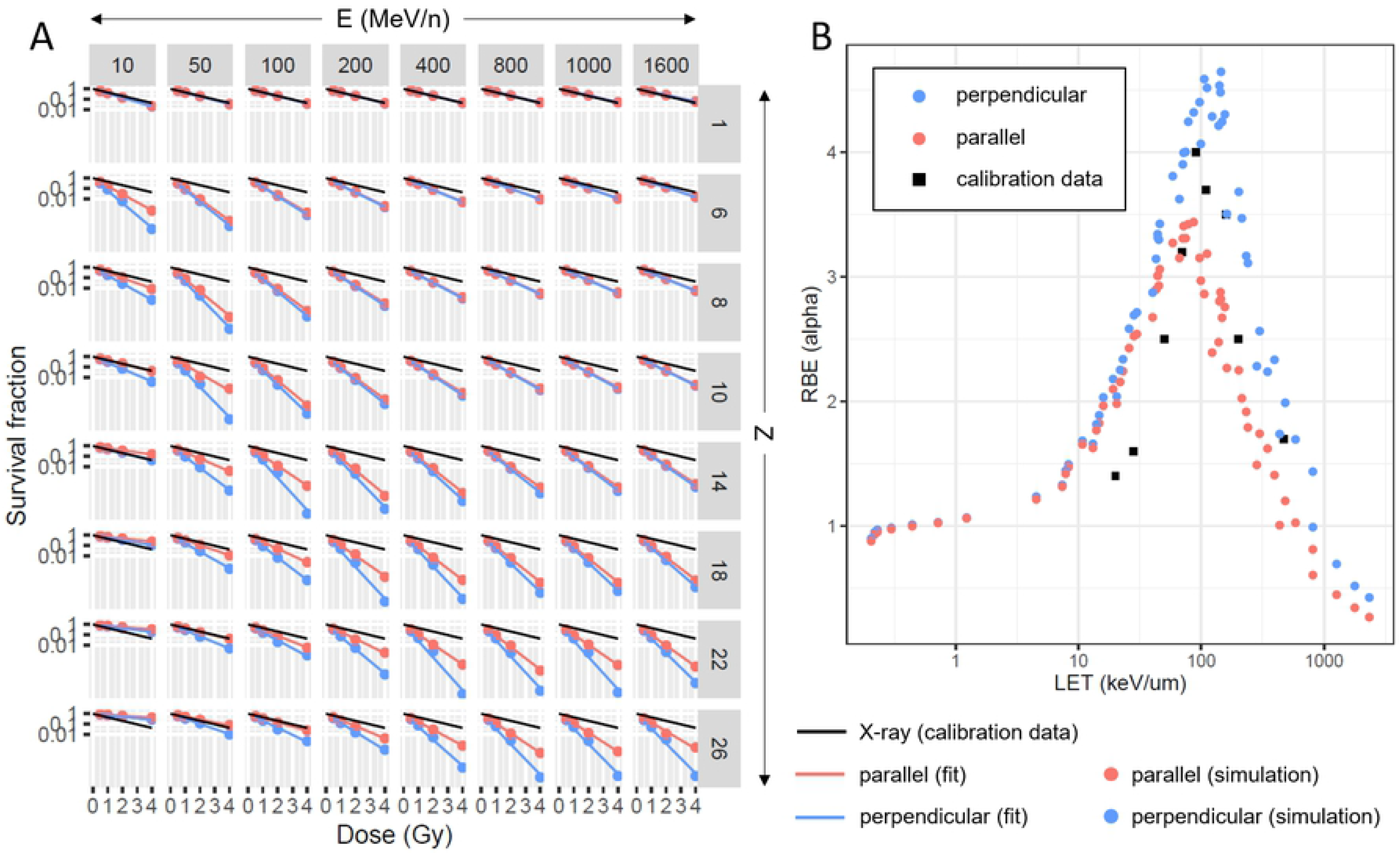
Cell survival predictions in HF19. A) Values obtained from our model for survival fraction in HF19 following exposure to ^2^H, ^4^He, ^16^C, ^18^O, ^20^Ne, ^28^Si, ^40^Ar, ^48^Ti, and ^56^Fe at E = 10, 50, 100, 200, 400, 800, 1000, 1600 MeV/n and D = 0.5, 1, 2, 4 Gy in perpendicular (blue) and parallel (red) orientations. The dots indicate direct simulation results based on our mathematical formalism using parameters from Figure 4. The lines indicate the corresponding fit, using the classic exponential model of survival as a function of dose: S = exp(-αD). The black line corresponds to experimental data reported for X-ray induced survival in HF19 [34]. B) RBE_α_ for cell survival in perpendicular (blue) and parallel (red) orientations. The RBE_α_ is computed as the ratio of the α parameter obtained from our simulations at each studied LET to the experimental α parameter reported for X-ray irradiation [34]. Black squares correspond to experimental values of RBE_α_ reported for a few LET [34].

**Figure 6.**
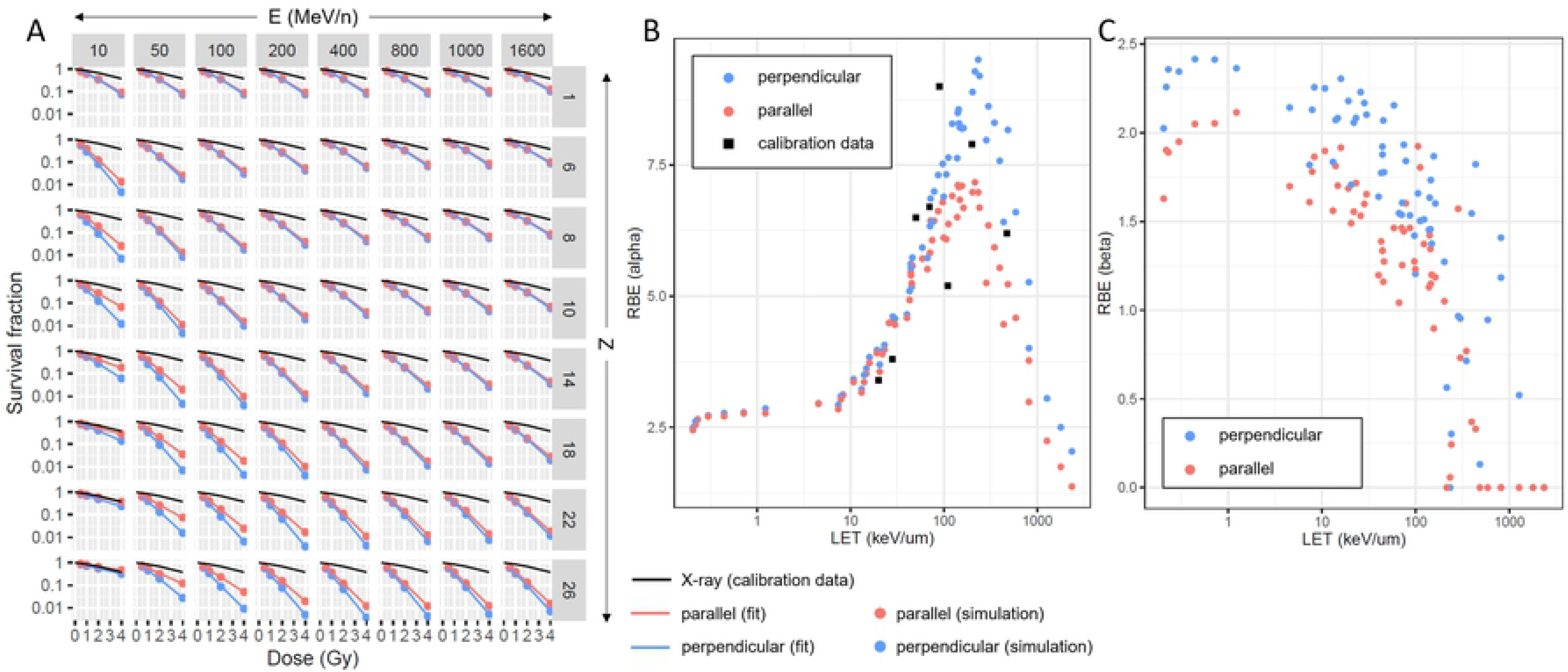
Cell survival predictions in V79. A) Values obtained from our model for survival fraction in V79 following exposure to ^2^H, ^4^He, ^16^C, ^18^O, ^20^Ne, ^28^Si, ^40^Ar, ^48^Ti, and ^56^Fe at E = 10, 50, 100, 200, 400, 800, 1000, 1600 MeV/n and D = 0.5, 1, 2, 4 Gy in perpendicular (blue) and parallel (red) orientations. The dots indicate direct simulation results based on our mathematical formalism using parameters from Figure 4. The lines indicate the corresponding fit, using the classic exponential model of survival as a function of dose: S = exp(-αD-βD^2^). The black line corresponds to experimental data reported for X-ray induced survival in V79 [34]. B) RBE_α_ and C) RBE_β_ for cell survival in perpendicular (blue) and parallel (red) orientations. The RBE_α_ (respectively RBE_β_) is computed as the ratio of the α (β) parameter obtained from our simulations at each studied LET to the experimental α (β) parameter reported for X-ray irradiation [34]. Black squares correspond to experimental values of RBE_α_ reported for a few LET [34].

**Figure 7.**
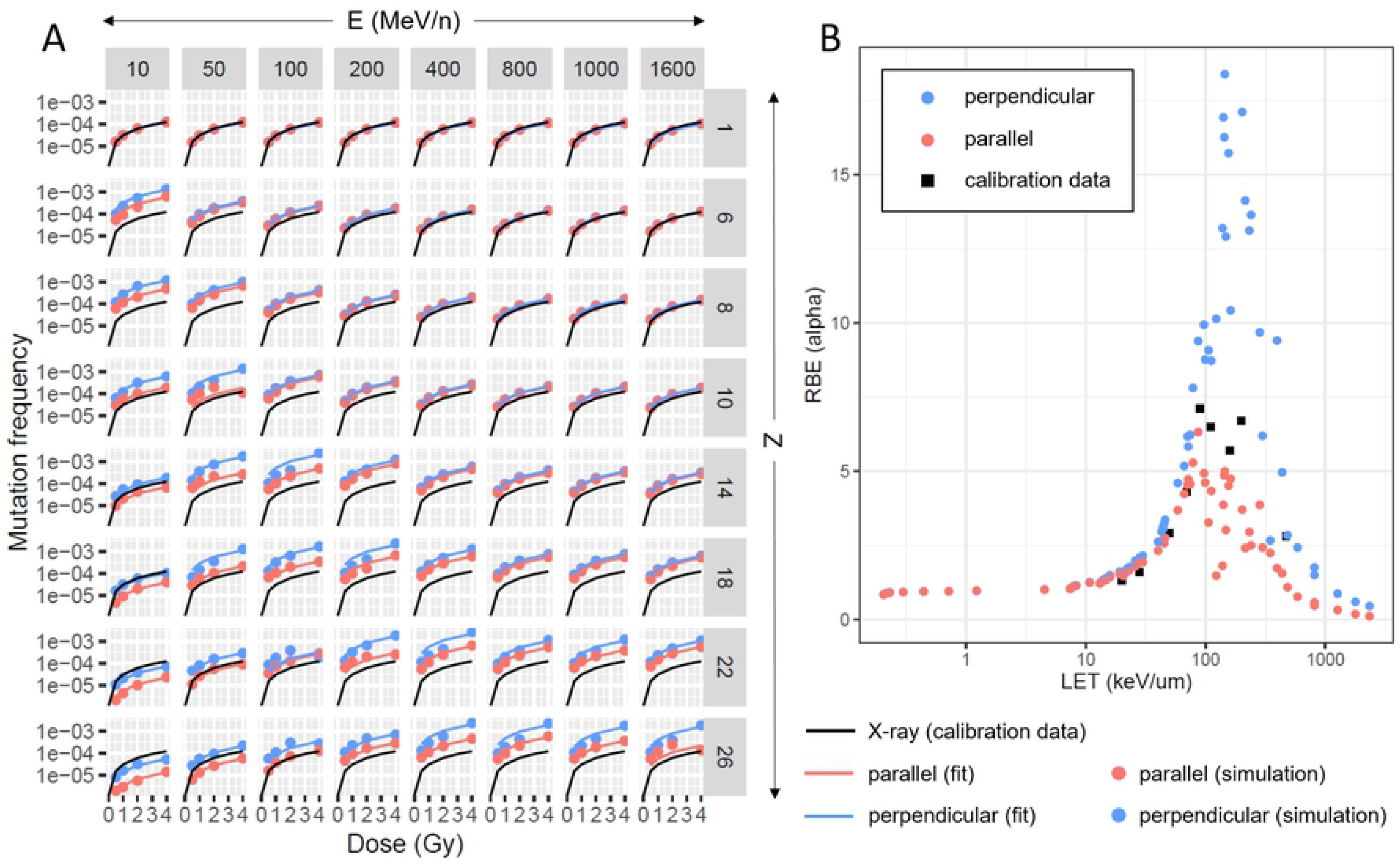
Mutation frequency predictions in HF19. A) Values obtained from our model for mutation frequency in HF19 following exposure to ^2^H, ^4^He, ^16^C, ^18^O, ^20^Ne, ^28^Si, ^40^Ar, ^48^Ti, and ^56^Fe at E = 10, 50, 100, 200, 400, 800, 1000, 1600 MeV/n and D = 0.5, 1, 2, 4 Gy in perpendicular (blue) and parallel (red) orientations. The dots indicate direct simulation results based on our mathematical formalism using parameters from Figure 4. The lines indicate the corresponding fit, using the classic linear model of mutation as a function of dose: M = αD. The black line corresponds to experimental data reported for X-ray induced mutation in HF19 [34]. B) RBE_α_ for mutation frequency in perpendicular (blue) and parallel (red) orientations. RBE_α_ is computed as the ratio of the α parameter obtained from our simulations at each studied LET to the experimental α parameter reported for X-ray irradiation [34]. Black squares correspond to experimental values of RBE_α_ reported for a few LET [34].

**Figure 8.**
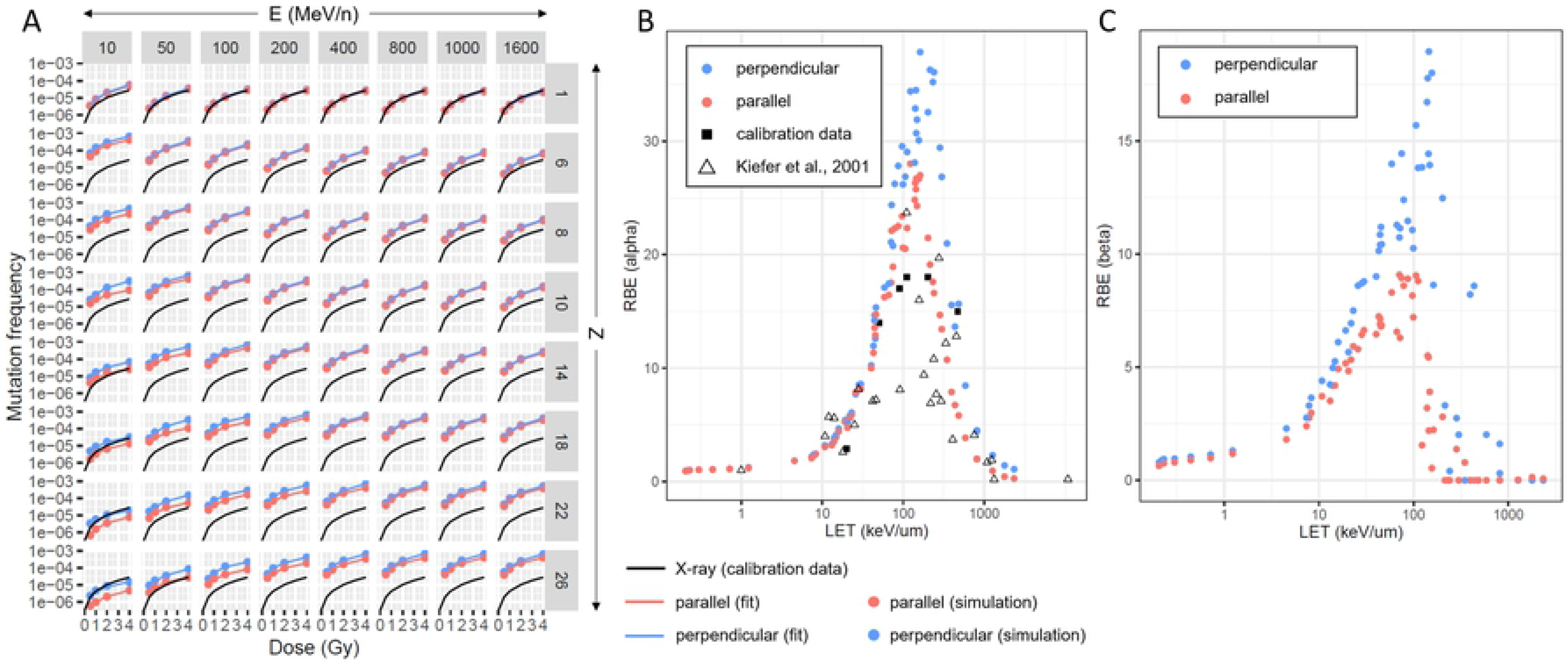
Mutation frequency predictions in V79. A) Values obtained from our model for mutation frequency in V79 following exposure to ^2^H, ^4^He, ^16^C, ^18^O, ^20^Ne, ^28^Si, ^40^Ar, ^48^Ti, and ^56^Fe at E = 10, 50, 100, 200, 400, 800, 1000, 1600 MeV/n and D = 0.5, 1, 2, 4 Gy in perpendicular (blue) and parallel (red) orientations. The dots indicate direct simulation results based on our mathematical formalism using parameters from Figure 4. The lines indicate the corresponding fit, using the classic quadratic model of mutation as a function of dose: M = αD + βD^2^. The black line corresponds to experimental data reported for X-ray induced mutation in V79 [34]. B) and C) RBE_α_ (respectively RBE_β_) for mutation frequency in perpendicular (blue) and parallel (red) orientations. RBE_α_ (RBE_β_) is computed as the ratio of the α (β) parameter obtained from our simulations at each studied LET to the experimental α (β) parameter reported for X-ray irradiation [34]. Black squares and grey triangles correspond to experimental values of RBE_α_ reported for a few LET in the paper used for the calibration of our model [34] and in another independent study [40].

Interestingly, for all LETs, both survival and mutation RBE_α_ are about twice as large for V79 (Figures 6B and 8B) compared to HF19 (Figures 5B and 7B), suggesting the higher radiation sensitivity of V79. In addition, we observe a clear effect of the beam orientation on RBE_α_ (and RBE_β_) for survival and mutation in HF19 (and V79), with higher RBE induced by perpendicular irradiation. The effect of beam orientation is stronger in HF19 compared to V79 for mutation RBE_α_, possibly because of different biological sensitivities, leading to different parameter fits. Finally, in V79, RBE_β_ for mutation drops quickly to 0 above 200 keV/μm, which indicates that the linear term dominates at this LET range.

In summary, in both cell types, survival and mutation RBE predictions using our model closely match the experiments used to calibrate the model parameters. In addition, to fully test the predictive power of our model, we also validated the modeled RBE against a large array of RBE values from other experiments using V79 cells that have been summarized by Kiefer et al. [40].

## DISCUSSION

In this study, we use the existence of DNA repair domains in mammalian cells to predict cell death and mutation frequency in 2 different cell types exposed to irradiation LET ranging from ~0.2 to ~2300 keV/μm, based on the coalescence into single RIFs of DSBs located in the same repair domain. Our team introduced a similar model in 2014 [28], which was only focused on predicting cell death in human breast cells, with limited confirmation in other cellular models. In this work, we introduce a formalism for mutation, test it in both human and hamster cell lines, assess its performance compared to experimentally obtained data, and evaluate the impact of beam orientation in a non-spherical cell geometry.

Both our model and the local effect model (LEM) [41] depart from the conventional older models of HZE track structure and induced cellular damage [42–46], where DSBs movements were considered minimum and passive with proximity-based clustering because of diffusion only [47, 48]. In contrast, our model is based on active movement of DSBs as suggested by recent literature [49–51], ignoring as a first approximation the complexity of individual DSB as being the main factor predictive of RBE. In this work, we show that the biological process of DSBs clustering within DNA repair domains is sufficient to explain and predict both cell death and mutation frequency, using the same formalism for both X-rays and HZE particles. In addition, unlike existing models of radiation-induced cellular damage, our experimentally-derived fitting parameters reflect a biological property of the DNA repair system instead of being abstract parameters. However, because of the observed LET dependence of the fitting parameters, prior calibration will be needed in the range of LETs of interest, which was not the case in our previous model limited to human breast cells to predict cell death only [28]. This suggests that the parameters of our model represent underlying LET-dependent biological effects, which also could be responsible for the higher sensitivity to the beam orientation observed in HF19 compared to V79 cells.

Our RBE predictions are in accordance with experimental results reported for HZE-induced cell death [37, 52, 53] and mutation frequency [25, 40], with an overkill effect above 200 keV/μm [54], and an RBE_β_ tending to zero for both survival and mutation at high LET. This RBE_β_ drop with increased LET corresponds to the linear dependence of the logarithmic survival probability and the mutation frequency with the irradiation dose for high-LET particles. Notably, our model was able to predict experimental data that were not used for calibration, reproducing the very high RBE previously observed in V79 for mutation [40].

Finally, our work highlights a biological prediction that rarely has been observed in the field of radiation biology, as it suggests that beam orientation may lead to distinct RBE in non-spherical cell geometry. Our study indicates higher cell death and mutation frequency when the irradiation beam is oriented perpendicular to the cell culture dish, compared to parallel orientation. It has been reported in the past that, in an ellipsoid cell, the number of chromosomal territories crossed by the HZE beam depends on the beam orientation [55]. Indeed, for a given irradiation ion, LET and dose, the number of particles is higher for an irradiation configuration orthogonal to the circular plane of the ellipsoid cell (perpendicular) compared to an irradiation along one of the large axes of the ellipse (parallel). While the deposition pattern is identical for both orientations in the direction of the beam, there is a higher proximity of DSBs between tracks in the perpendicular orientation for a given dose, because of the larger number of particles. This could be responsible for more DSBs combinations in perpendicular irradiation, and subsequent increase in cell death and mutation frequency, which remains to be tested experimentally.

In summary, our results show that the DSBs distribution in the cell is sufficient to predict cellular consequences of HZE particles irradiation and we demonstrate that the orientation of asymmetric cells relative to the beam can modify biological responses to radiation. Such prediction, if confirmed experimentally, would be another strong supporting evidence for DNA repair domains and their critical role in interpreting cosmic radiation sensitivity. It also would have very important consequences in hadron therapy where cell shape and orientation may have to be taken into account to better predict the response of healthy surrounding tissues. This is especially true for tissue with strong cell orientation like the brain cells. In the context of space exploration and prediction of GCR-induced cancer risks, the orientation of the cell relative to the irradiation incidence appears to be a determinant factor to RBE values, which should be further investigated in the future to improve prediction reliability. In particular, we anticipate from our results that corrections will be required to translate *in vitro* data in plated cells to *in vivo* risk assessment in cells presenting their physiologic conformation in tissues.

While the true nuclear organization of repair domains remains to be discovered and characterized, we have shown in the past that these domains are located most likely in the euchromatin and that DSBs formed in the heterochromatin need to be moved by Rad51 to the euchromatin to complete repair [49, 56], with displacement magnitudes on the order of 1 to 2 μm [50]. Cubical domains of identical size are therefore a clear over-simplification and there would be value to improve this work by modeling the chromatin organization and the active mechanism moving DSBs in the nucleus. This work already uses a sophisticated representation of the nucleus, with each chromosome and chromatin loop independently modeled. It would therefore be interesting to push this work further to model chromosomal rearrangements following ionizing radiation, an important hallmark of cancer.

## MATERIALS AND METHODS

### DSBs simulations

The DSBs distributions induced by radiation tracks in ellipsoid cells were generated using the simulation code RITCARD [29]. This model brings 3 entities together: 1. a model for the chromosome structures, based on random walk with monomers of 20 nm, with the addition of chromatin loops and chromosomal domains, 2. a model for the stochastic track structure, which calculates the particle transport and energy deposition in voxels of 20 nm [57], corresponding to the size of the monomers of the chromosome model, and 3. a model for the intersections between voxels with dose greater than 0, and the corresponding chromosome monomers. The number of DSBs in a monomer is determined by sampling the Poisson distribution, 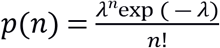, where λ=QD, Q is the sensitivity parameter corresponding to a constant DSBs yield (α), and D is the dose to the voxel/monomer. Alpha is assumed to be 35 DSB/Gy for high LET and 25 for low LET [58]. One cell geometry was considered in this work (Figure 1B), with the nucleus being a 3-dimensional ellipsoid (a = b = 17 μm and c = 6 μm). We generated N_cell_ = 1000 nuclei for each of the 576 exposure conditions: particles range: Z = 1, 2, 6, 8, 10, 14, 18, 22, 26 (respectively: H, He, C, O, Ne, Si, Ar, Ti, Fe); energy range: E = 10, 50, 100, 200, 400, 800, 1000, 1600 MeV/n, dose range: D = 0.5, 1, 2, 4 Gy, and beam orientation: perpendicular and parallel. This led to a large coverage of LETs from ~0.2 keV/μm to ~2300 keV/μm. Note that for each simulated dose, all 1000 simulated nucleus received a different individual dose, averaging to the desired overall dose and obeying the expected Poisson distribution in terms of direct particle traversal per cell.

### Model fitting

Model fitting was performed using Prism and nlme (nonlinear mixed-effects) R package for linear and nonlinear mixed effects models [59].

## ACKNOWLEDGEMENTS

This work was also originally supported by the Lawrence Berkeley National Laboratory Innovation grant #DE-AC02-05CH11231 and NASA grant NNL15AA08I. E.P. is funded by the Translational Research Institute for Space Health through NASA Cooperative Agreement NNX16AO69A. S.B. was supported by the Human Research Program of the Human Exploration and Operations Mission Directorate of the National Aeronautics and Space Administration. I.P. and A.P were supported by the NASA Human Health and Performance contract NNJI5HK11B We thank Dr. Cekanaviciute at NASA Ames Research Center and Dr. Floriane Poignant at NASA Langley Research Center, for the detailed feedback on this manuscript.

## SUPPORTING INFORMATION CAPTION

**Figure S1. Calibration of the parameters of the model.** Experimental data (black points) for probability of survival and mutation frequency in HF19 and V79 at each tested LET [34], and corresponding fits with optimal parameters (*γ_1_* and *γ_2_* for survival, *δ_1_* and *δ_2_* for mutation) for each domain size (colored lines).

